# Gonadotrophs have a dual origin, with most derived from pituitary stem cells during minipuberty

**DOI:** 10.1101/2024.09.09.610834

**Authors:** Daniel Sheridan, Probir Chakravarty, Gil Golan, Yolanda Shiakola, Jessica Olsen, Elise Burnett, Christophe Galichet, Patrice Mollard, Philippa Melamed, Robin Lovell-Badge, Karine Rizzoti

## Abstract

Gonadotrophs are the essential pituitary endocrine cells for reproduction. They produce both luteinizing (LH) and follicle-stimulating (FSH) hormones that act on the gonads. Gonadotrophs first appear in the embryonic pituitary, along with other endocrine cell types, and all expand after birth. We show here that most gonadotrophs originate from a population of postnatal pituitary stem cells during minipuberty, while those generated in the embryo are maintained, revealing an unsuspected dual origin of the adult population. This has implications for our understanding of the establishment and regulation of reproductive functions, both in health and in disease.

## Introduction

The reproductive axis comprises the hypothalamus, the pituitary gland and the gonads. While all its components are assembled in the embryo, it only starts to be active postnatally, initially during a transient period known as minipuberty, which is important for future male fertility and cognitive development in both sexes (*1*). Activity will start again at puberty, which marks the onset of reproductive capacity. The hypothalamic gonadotrophin releasing hormone (GnRH) has a central regulatory role during these periods (*2*). Its pulsatile secretory patterns are regulated by a neuronal network that integrates peripheral signals; it is released in the hypophyseal portal system, through which it reaches the pituitary where it controls production and secretion of luteinizing (LH) and follicle stimulating (FSH) hormones. The two gonadotrophins act in turn on the gonads, regulating steroid and gamete production. Gonadal steroids exert crucial feedback both at the hypothalamic and pituitary level.

Most adult pituitary gonadotrophs produce both LH and FSH. However, the patterns of their secretion differ; LH secretion closely follows GnRH pulses, while FSH does not, and it is regulated by additional peptide hormones. The modalities of LH and FSH regulation are not completely understood, in particular how secretion of LH and FSH is differentially regulated by the same ligand (*3*). Adding to this complexity, we reveal here that gonadotrophs comprise a dual population based on different developmental origins.

Gonadotrophs initially arise in the embryonic pituitary. They are amongst the first endocrine cell types to commit, at 12.5dpc in the ventral pituitary primordium, with upregulation of the glycoprotein α, the subunit common to LH, FSH and TSH (thyroid stimulating hormone). LHβ and FSHβ subunits are only expressed from 16.5dpc in the mouse, which is considered the true gonadotroph birthdate, and GnRH is involved in this, at least in males (*4*). Postnatally, gonadotrophs expand dorsally thorough the gland as a second wave during the first two weeks after birth (*5-7*).

We and others have shown that the pituitary contains a population of stem cells (SCs) (*8*). These do not play a significant role during normal turnover in the adult gland. However, their contribution to the gland’s rapid growth postnatally could not be defined because of inefficient genetic lineage tracing tools and use of tamoxifen, a selective estrogen receptor modulator which perturbs normal physiology (*9, 10*). Here, we performed lineage tracing using a new, efficient and more physiologically neutral doxycycline-dependant *Sox2^rtTA^* lineage tracing tool and uncovered that the majority of gonadotrophs differentiate in both sexes from postnatal pituitary SCs, during a period encompassing minipuberty. However, their differentiation does not depend on GnRH or gonadal feedback. These gonadotrophs invade the gland from the SC niche, with the exception of a small ventral domain where embryonic gonadotrophs remain confined. The discovery of a dual origin for gonadotrophs may help understand aspects of gonadotrophin regulation and mechanisms of diseases affecting puberty and fertility.

## Results

### Endocrine cell type-specific mechanisms underline postnatal expansion

In mouse and humans, all the 6 different pituitary endocrine lineages, somatotrophs secreting growth hormone (GH), lactotrophs secreting prolactin (PRL), thyrotrophs secreting TSH, corticotrophs secreting Adrenocorticotrophic hormone (ACTH), melanotrophs secreting melanotroph stimulating hormone (MSH) and gonadotrophs, emerge in the embryonic pituitary (*11, 12*). In the mouse, this is followed postnatally by a period of significant growth until the gland reaches its mature size (*13, 14*), following both an increase in cell numbers, due to proliferation of both stem and endocrine cells (*15-17*), and endocrine cell swelling, as the secretory apparatus matures (*18*). Along with SCs, a population of POU1F1^+ve^ hormone^-ve^ progenitors, committed to lactotroph, somatotroph or thyrotroph fate, has also been reported (*17*). To better characterise this phase, we examined how proportions of each endocrine population evolve, by performing automated cell counting on dispersed pituitary cells stained for the hormone they secrete, from P5 to one-year-old in both males and females (Fig.1A, Table S1). In adults, the percentage of endocrine cells obtained for each population (Fig.1A) match what had been described previously (*19*). From P5 to adulthood, the proportion of somatotrophs and lactotrophs increased the most, leading to their known sex-specific dominance in the male and female pituitaries, respectively (Fig.1A). Gonadotrophs also exhibit significant postnatal growth during the three first postnatal weeks. In contrast, the proportions of corticotrophs and thyrotrophs, which are near their highest at birth, either remain stable or decrease as the gland matures. To better visualize how each population evolves, we related cell-type proportions to their initial percentage at P5 (Fig.1B). In both sexes lactotrophs prominently increase in agreement with their mainly postnatal expansion. Representing the second largest increase, both gonadotrophs and somatotrophs display a similar pattern of expansion, undergoing an approximately 3-fold increase in proportion.

**Figure 1.**
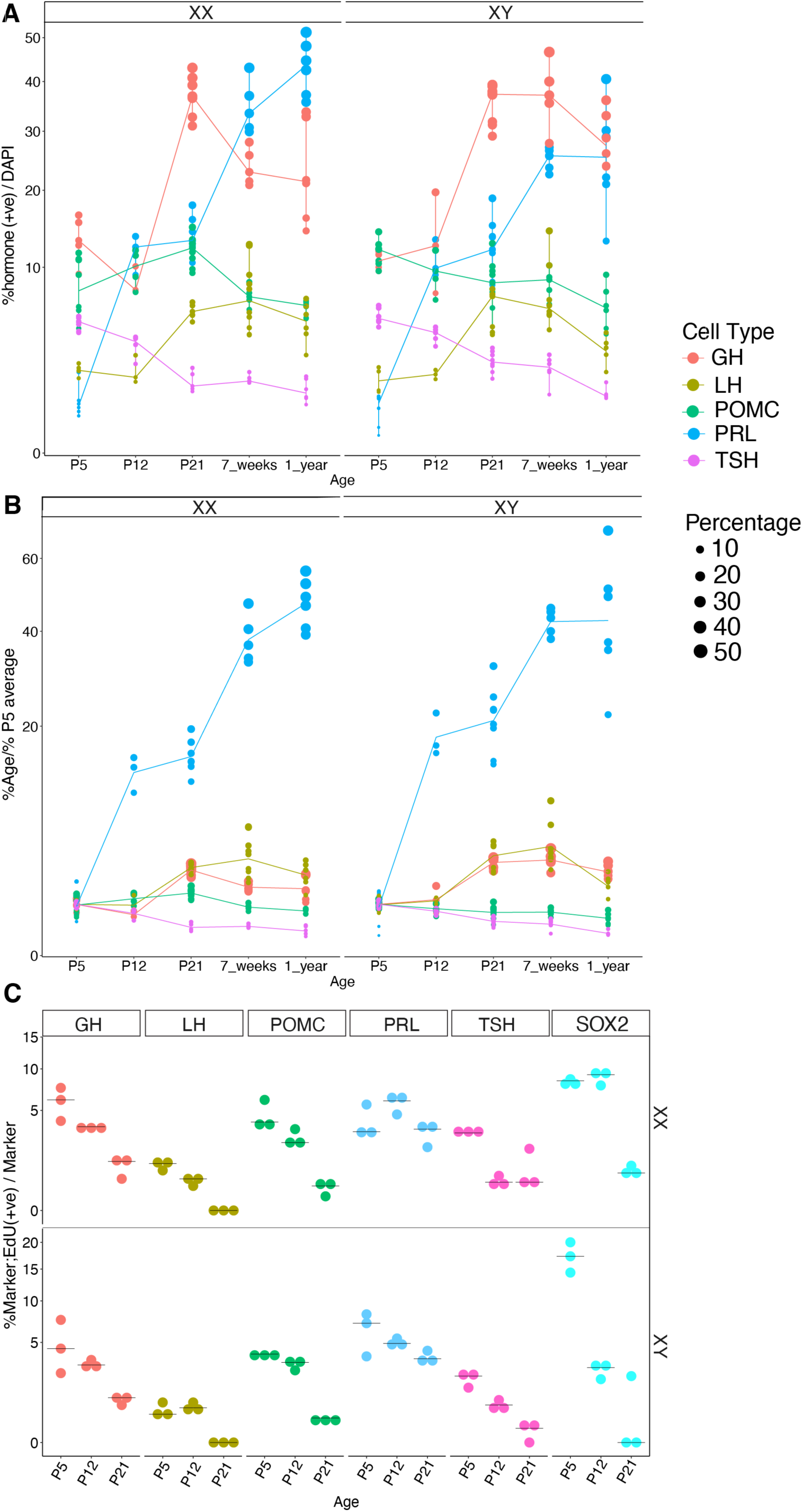
Cell type-specific expansion in the postnatal pituitary. **A)** Percentages of endocrine cells from dissociated pituitaries stained for each endocrine hormone (GH for somatotrophs, PRL for lactotrophs, TSH for thyrotrophs, LH for gonadotrophs and POMC for both corticotrophs and melanotrophs) from P5 to one-year old males and females. **B)** Evolution of each population: the percentage at each age is shown in relation to the average percentage at P5. The size of dots relates to the proportion of each population in the gland. **C)** Percentage of EdU positive cells/marker following a one-hour EdU pulse. Each dot represents one pituitary.

To investigate mechanisms underlying populations expansion, we assessed cell proliferation during the first three weeks post-birth by performing one-hour EdU pulses (Fig.1C, Fig.S1, Table S2). From P5 to P21, somatotrophs, lactotrophs and POU1F1^+ve^;GH/PRL/TSH^-ve^ progenitors had the highest percentage of proliferative cells, which correlates with the expansion of these cell types and previous observations (*15-17*). In contrast, gonadotrophs, despite the remarkable increase in their population size, had the lowest proliferation rate of all endocrine types at all assessed time points. Finally, SOX2^+ve^ SCs showed high proliferation rates early on, with significantly more proliferation in males at P5 (P<0.05, Table S2), which then declined noticeably between P12 and P21, as reported (*15, 16*).

These results expand previous analyses, highlighting the differential growth of each endocrine population, and show that different mechanisms underlie their evolution. While somatotrophs and gonadotrophs appear to progress at a comparable rate, their different proliferative indices suggest that although division of existing somatotrophs may play a role in their expansion, this is not the case for gonadotrophs.

### Neonate SOX9iresGFP^+ve^ SCs are predicted to contribute to post-natal pituitary endocrine cell emergence

To characterise the SCs during their highly proliferative period and assess their contribution to the formation of new endocrine cells, we performed single cell RNAseq on the SOX9iresGFP^+ve^ fraction at postnatal day 3 (P3) in males and females. This fraction comprises SCs and their immediate progeny because GFP persists longer than SOX9, allowing for a short-term tracing of the SCs (*20*). Male and female datasets were integrated (Fig.2A, Fig.S2). Uniform Manifold Approximation and Projection (UMAP) allowed clustering of *Ednrb*^+ve^ and *Aldh1a2*^+ve^ SCs, representing respectively cleft and anterior lobe SCs (*20*). Clusters corresponding to all endocrine cell types were present in the dataset, with gonadotrophs representing the largest endocrine cluster. We also observed a *Pitx1*^-ve^ *Decorin*^+ve^ cluster labelled “extra-pituitary origin” (Fig.S2); this may contain mural cells of neural crest origin, as we showed in adults (*20*). We observe a relatively high proportion of proliferative cells, as expected (Fig.1C) (*15*), and the presence of two clusters in which expression of SC markers decreases, but are negative for any endocrine marker, suggesting that they represent differentiating SCs (thus named SC differentiating 1 and 2). These clusters are characterised by expression of the WNT pathway transcriptional activator *Lef1*, suggesting activity of the pathway, which is consistent with the proposed paracrine role of SC-derived WNT on their differentiating progeny (*21*). However, the expression of cell cycle inhibitors *Cdkn1a* and *Gadd45g* (Fig.S2) suggests that cell proliferation is limited in these progenitors.

**Figure 2.**
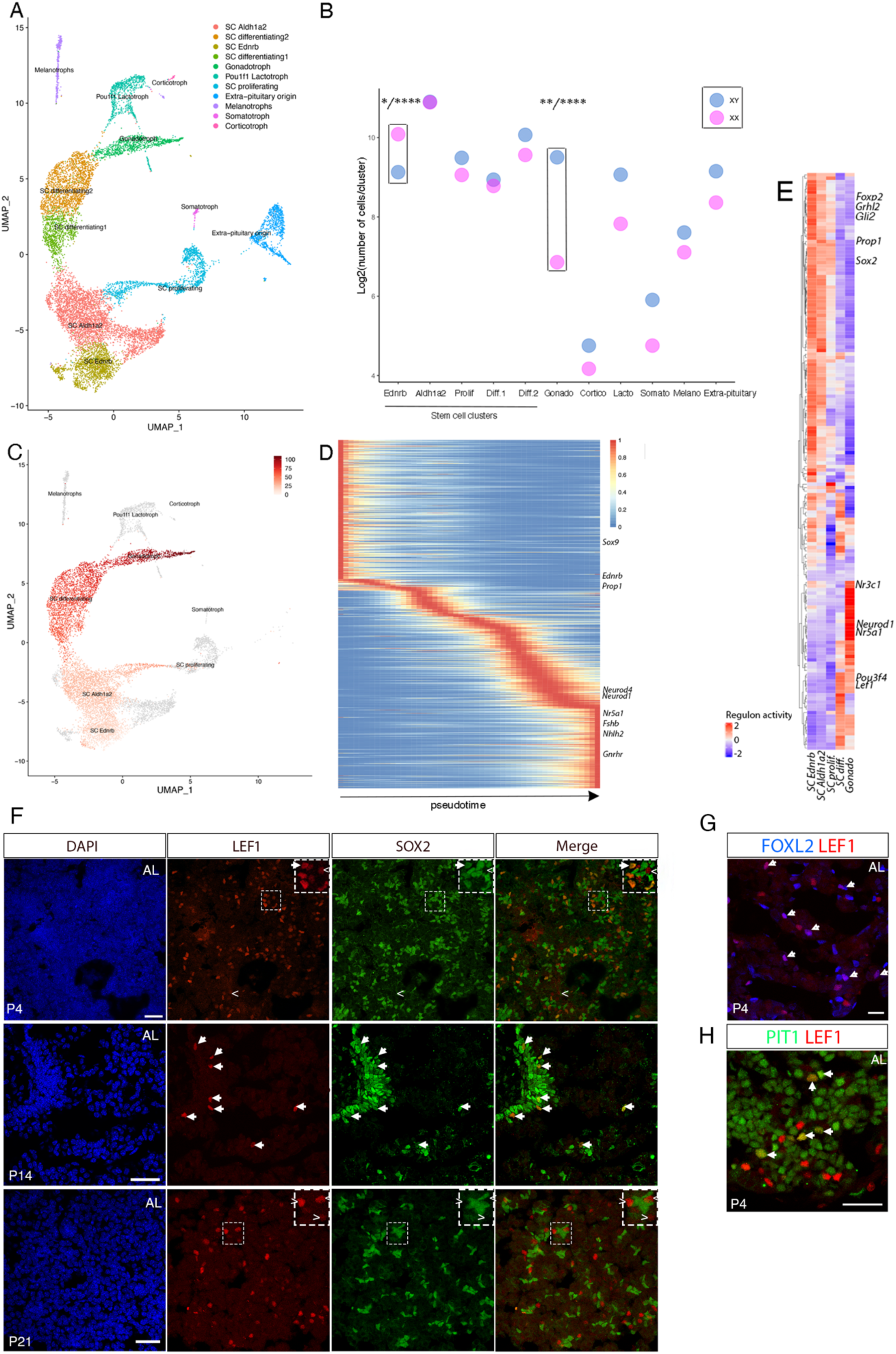
Single cell analysis of early postnatal stem cells and their immediate progeny. **A)** UMAP clustering of P3 male and female SOX9iresGFP^+ve^ cells. **B)** Pair-wise comparison for proportion test performed on male and female cells. Significance is exclusively shown for clusters where distribution was different from all other clusters. This shows that the proportion of Ednrb^+ve^ (cleft) SCs is superior in females while the proportion of differentiating gonadotrophs is higher in males. **C)** UMAP representation of the pseudotime gonadotroph trajectory from Ednrb^+ve^ SCs to gonadotrophs. Extra-pituitary cells (Pixt1^-ve,^ Sup. Fig.2) were filtered out. **D)** Heatmap showing genes whose expression pattern correlate with the gonadotroph pseudotime trajectory. **E)** Heatmap displaying regulons according to the gonadotroph trajectory pseudotime. **F)** Double immunostaining for LEF1 and SOX2 from P4 to P21 in male pituitaries. At P4, LEF1 is mostly co-expressed with SOX2 (arrow), with a small number of SOX2^-ve^;LEF1^+ve^ cells nearby (indicated by <). At P14, LEF1 remains co-expressed with SOX2. By P21, LEF1 no longer colocalises with SOX2. **G**, **H**) At P4, LEF1 is co-expressed in some FOXL2^+ve^ (G) and POU1F1^+ve^ (H) progenitors. Scale bars represent 30 μm in all panels.

We next looked for sex differences in cluster contribution (Fig.2B). A pairwise comparison for proportion test showed that the gonadotroph cluster comprised more male cells and conversely, that the *Ednrb*^+ve^ SC cluster had more female cells. This suggests that male SCs differentiate more into gonadotrophs than female SCs at this stage.

Pseudotime analyses were performed using Slingshot (*22 (Fig.2C, D)*), and gene regulatory networks (GRN) analysis using SCENIC (*23*) (Fig.2E). The SCENIC analysis identifies regulons consisting of a transcription factor and its putative targets and quantifies their activity according to the expression patterns. Trajectories toward all endocrine cell types were projected (Fig.2C, Fig.S3) with a common root starting from *Ednrb*^+ve^ SC then progressing to *Lef1*^+ve^ clusters. Given that it was the predominant derivative endocrine cell type, we focussed our attention on the gonadotroph trajectory. These analyses showed known regulators and markers of gonadotroph fate acquisition (*Nr5a1*, *Fshb*, *Gnrhr*, Fig.2D, E), but also transcripts from genes encoding the bHLH transcription factors *Neurod1*, *4* and *Nhlh2* suggesting that these are likely involved in gonadotroph differentiation as well. In the embryo, *Ascl1*;*Neurod1/4* deleted embryonic pituitaries show reduced gonadotroph numbers (*24*) while puberty is impaired in *Nhlh2^-/-^* animals (*25*). In the latter, the number of GNRH neurons is reduced but the pituitary is also affected. Our analyses suggest that these factors are involved during terminal differentiation of gonadotrophs. In addition, our data show that *Foxp2*, a marker of gonadotrophs in the adult (*26*), is expressed in SCs and progenitors. Its presence in all lineages in our SCENIC analyses, as previously seen in adult pituitaries (*27*), implies an involvement in endocrine cell fate acquisition (Fig.2E, Fig.S3). Similarly, the presence of *Gli2* in all our GRN analyses suggest that the Hedgehog (Hh) pathway is active in differentiating SCs. The pathway is important for pituitary embryonic development and potentially involved in adult pituitary tumor formation (*28*). We find that the only Hh pathway ligand to be expressed in our dataset is *Shh*, exclusively present in SC differentiating 2 (Fig.S2), while the Shh receptor, Ptch1, is also expressed in the SCs. This suggests that differentiating cells interact with more naïve SCs, where the *Gli2* regulon is predicted to be active. In agreement with a role for the WNT signalling pathway, a *Lef1* GRN is predicted to be involved in all trajectories. LEF1 protein expression pattern is very dynamic during the first three postnatal weeks. At P4 we mostly see SOX2;LEF1 double^+ve^ cells, and our scRNAseq analyses suggest that proliferation is reduced in these cells compared to more naïve SOX2^+ve^;LEF1^-ve^ SCs (Fig.2F). We also see LEF1 co-localisation with POU1F1 or FOXL2 (Fig.2G,H), correlating with its expression in a transient differentiating population, likely giving rise to lactotrophs and gonadotrophs respectively, according to the UMAP (Fig.2A and Fig.S2). In contrast, from P21, LEF1 marks a distinct, SOX2-negative cell population, often in close proximity to SOX2^+ve^ cells (Fig.2F), as previously shown for its transcript, and these have been shown to be highly proliferative (*21*), at a time where in turn, proliferation of SOX2^+ve^ cells diminishes (Fig.1C).

All together, these data suggest that SCs mostly give rise to gonadotrophs at P3. They predict known and novel regulators during this period of high SC activity, with a likely involvement for WNT and SHH signalling.

### SOX2^+ve^ postnatal SCs give rise to most adult gonadotrophs

To examine and expand our transcriptomic analysis predictions, we performed lineage tracing analyses. We generated a *Sox2^2A-rtTA^* allele where both copies of *Sox2* are maintained in order to avoid the hypopituitarism seen in *Sox2^+/-^* animals (*29*) and the use of tamoxifen. We generated *Sox2^2A-rtTA/+^;TetO-Cre;Rosa26^ReYFP/+^*(*Sox2rtTA;eYFP*) animals and induced recombination with two 9TBDox injections (*30*), at P0 and P1. 24 hours after the last treatment, approximately 70% of SOX2^+ve^ cells were traced, with all labelled cells expressing SOX2, demonstrating efficiency and specificity of the system (Fig.S4). Moreover, treated pups gave rise to fertile adults with normal GH and LH levels (Fig.S4).

We then analysed the progeny of SCs in *Sox2rtTA;eYFP* pituitaries induced at birth, from P5 and up to one year (Fig.3A). We observe eYFP;hormone double^+ve^ cells for all lineages, as shown previously (*9, 10*). We quantified SC post-natal contribution to each anterior lobe endocrine population from P5 to one-year old, in both sexes (Fig.3B, C, Table S3). At all ages examined, postnatal SCs contribute very little to somatotroph and thyrotroph populations, both below 1%. A higher proportion of SC progeny is noted in lactotrophs and corticotrophs, ranging from 7 to 12%. Remarkably, comprising up to 80% of eYFP^+ve^ cells, we find that gonadotrophs are the population to which SCs contribute by far the most. These results were similar in both sexes.

**Figure 3.**
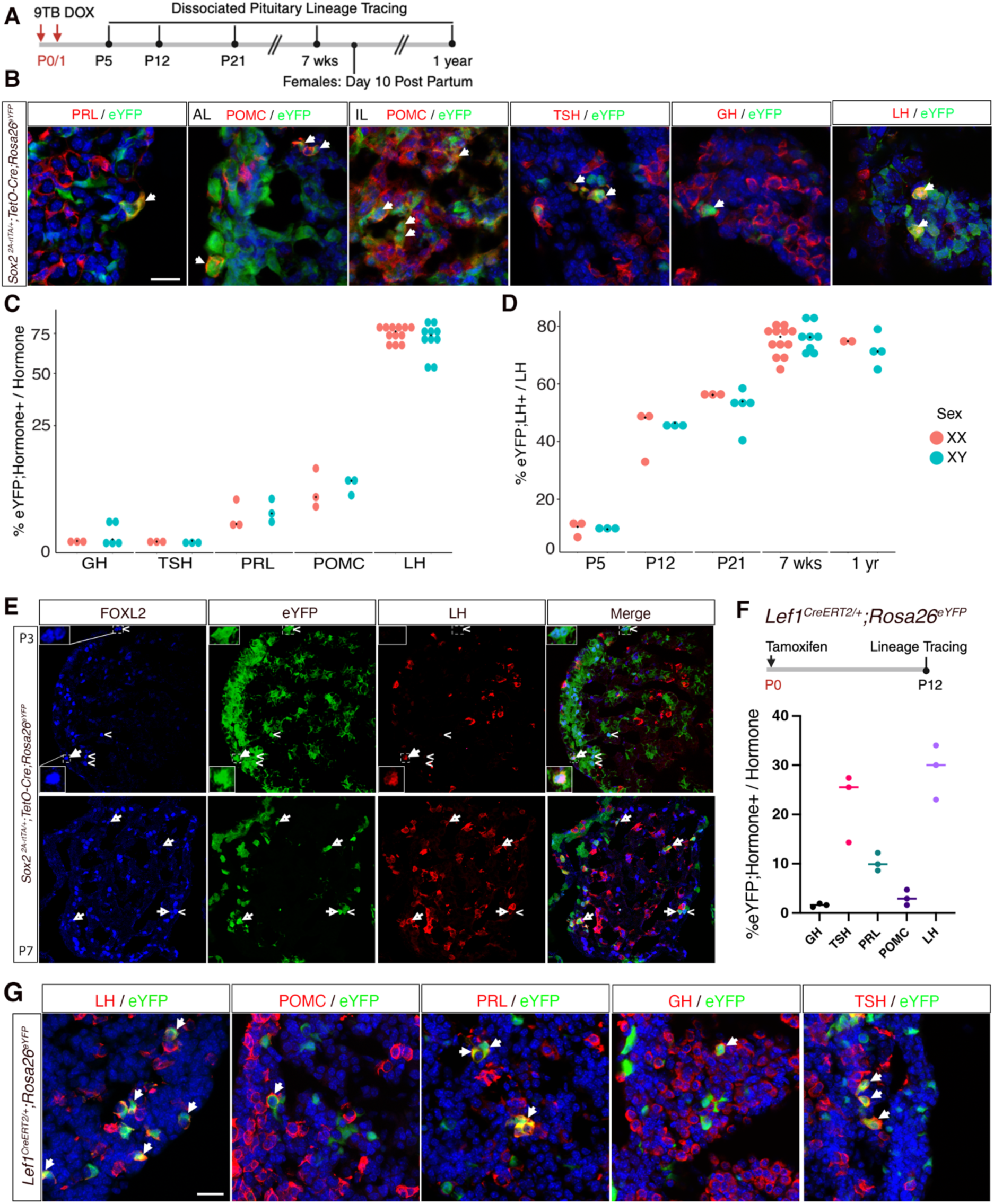
SOX2^+ve^ SC and LEF1^+ve^ progenitors give rise to most adult gonadotrophs during a period encompassing minipuberty. **A)** Timeline of postnatal lineage tracing induction in *Sox2^2A-rtTA/+^;TetO-Cre;Rosa26^ReYFP/+^* and harvest timepoints for quantification. **B)** SCs contribute to all endocrine lineages, as seen by eYFP co-localisation with each hormone in a 7-week-old male. **C)** Quantification of SC contribution to each cell type in both sexes in 7-week-old mice, as measured by the percentage of eYFP;hormone double^+ve^ cells in the total hormone^+ve^ population. **D)** Time course of postnatal SC contribution to the gonadotroph population from P5 to 1 year of age. There is no difference between sexes in SC contribution to any cell type (P>0.05, multiple unpaired t-test with Benjamini-Hochberg post hoc test). **E)** SCs rapidly commit to the gonadotroph lineage postnatally, with numerous eYFP;FOXL2 double^+ve^;LH^-ve^ cells present at P3 (top), (<). From P7 onwards (bottom), most eYFP;FOXL2 double^+ve^ cells are also LH^+ve^, (arrow). **F)** Timeline of lineage tracing induction in *Lef1^CreERT2/+^;Rosa26^eYFP^*^/+^ mice, with tissue counts performed at P12. **G**) LEF1^+ve^ cells contribute to all endocrine cell types, as evidenced by the co-localisation of eYFP with all hormones in a P12 male. Scale bar = 20 µm for all panels. Graphs show individual data points (each dot represent one animal) and group median. AL: anterior lobe, IL: intermediate lobe.

We then analysed the temporal dynamics of SC-derived endocrine cell emergence (Fig. 3C, Fig.S5). For gonadotrophs, from the 10% of the small number of LH;FSH double^+ve^ cells present at P5, eYFP^+ve^ cell contributions rise sharply to over 40% by P12, surpassing 50% by P21 (pre-weaning), reaching and stabilising at approximately 75% by 7 weeks and up to one year (Figure 3D and Table S4). To further detail the tempo of gonadotroph fate acquisition, we looked at expression of the transcription factor FOXL2, a marker of gonadotroph and thyrotroph commitment (*31*) (Fig.3E). At P5, most SOX2^-ve^;eYFP^+ve^ cells are already positive for FOXL2, while negative for both gonadotroph and thyrotroph hormones, suggesting that cell fate commitment occurs soon after birth. Furthermore, we observe concomitant up-regulation of both LH and FSH from early stages of differentiation in newly differentiated gonadotrophs (Fig.S6), while the UMAP suggests a delayed or reduced expression of *Fshb* compared to *Lhb* in differentiating gonadotrophs (Fig.S2). To validate our characterisation of LEF1 as a marker of a transient differentiating population, we performed lineage-tracing using *Lef1^CreERT2/+^;Rosa26^eYFP^*^/+^ mice, inducing at P0 (Fig.3F, G). In agreement with our hypothesis, LEF1 progeny contributed to all anterior lobe endocrine cell types (Fig.3E). Quantification at P12 revealed a proportion of eYFP^+ve^ gonadotrophs similar to that found using *Sox2rtTA;eYFP* (Fig. 3F). There was a noticeable increase in eYFP^+ve^ somatotrophs and lactotrophs compared to the SOX2 tracing, potentially due to LEF1 being expressed in the POU1F1 lineage, or to the effect of tamoxifen. Additionally, Lef1 is expressed in thyrotrophs (using the dataset published in (*32*)), explaining their presence in our quantification.

Finally, to explore the effects of physiological changes during pregnancy, we analysed the SC contribution to all endocrine cell types in lactating dams and age-matched virgin females. No significant change was observed (Fig.S7).

In conclusion, most adult gonadotrophs originate from postnatal SCs during a period encompassing minipuberty (*33*). This result fits with the low proliferation rate of gonadotrophs (Fig.1C) as their expansion is explained by the differentiation of SC, as we have shown. It is notable that the SCs contribute little to other endocrine populations, and pregnancy does not alter patterns of differentiation. WNT signalling is likely to be important during cell fate acquisition since all committed progenitors descend from LEF1^+ve^ progenitors. Finally, gonadotroph cell fate acquisition occurs rapidly, with a significant proportion of *Sox2rtTA;eYFP* labelled progeny expressing FOXL2 already at P5.

### Embryonic and postnatal-born gonadotrophs are located in distinct domains

Examination of traced *Sox2rtTA;eYFP* pituitaries at P3 revealed a small, eYFP^-ve^, LH^+ve^ population confined to the medio-ventral surface of the pituitary (Fig. 4A, upper panel). In contrast, at P15, eYFP^+ve^ gonadotrophs are enriched dorsally and seem to emerge from the SCs lining the cleft, while the eYFP^-ve^;LH^+ve^ population remains confined ventrally (Fig. 4A, lower panel). Several eYFP^-ve^ gonadotrophs are present away from the ventral medial region; because the tracing method is not 100% efficient (Sup.Fig.4), these are likely to represent non-recombined SC progeny. To quantify the preferential localisation of eYFP^+ve^ gonadotrophs, we sectioned the entire pituitary transversely, designating the initial 50% of sections as ventral and the latter half as dorsal (Fig. 4B). By P7, corresponding to an early stage of SC differentiation, the prevalence of LH;eYFP double^+ve^ gonadotrophs was comparably low (<10% of LH^+ve^ cells) across both regions. In contrast, by P21, close to 80% of dorsal gonadotrophs were eYFP^+ve^, while only approximately 30% of LH^+ve^ cells in the ventral region were derived from SCs. This disparity underscores the substantial postnatal differentiation of SCs to gonadotrophs within the dorsal region, predominantly emerging from, or adjacent to the pituitary cleft. This finding correlates well with the pseudotime trajectory, which predicts an *Ednrb^+ve^*, therefore cleft SC origin of gonadotrophs (Fig.2C) (*20*). To investigate this, we generated an *Ednrb^2ArtTA^* allele to exclusively trace cleft SCs. In *Ednrb^2A-rtTA/+^;TetO-Cre;Rosa26^ReYF/+^*animals induced at birth, we observe a robust contribution of the progeny to the gonadotroph lineage, demonstrating at least a partial cleft-lining origin of SC-derived gonadotrophs (Sup.Fig8). These may invade the pituitary later; conversely, more ventrally located parenchymal SCs may undergo differentiation, explaining the presence of SC-derived gonadotrophs in the ventral half of juvenile glands.

**Figure 4.**
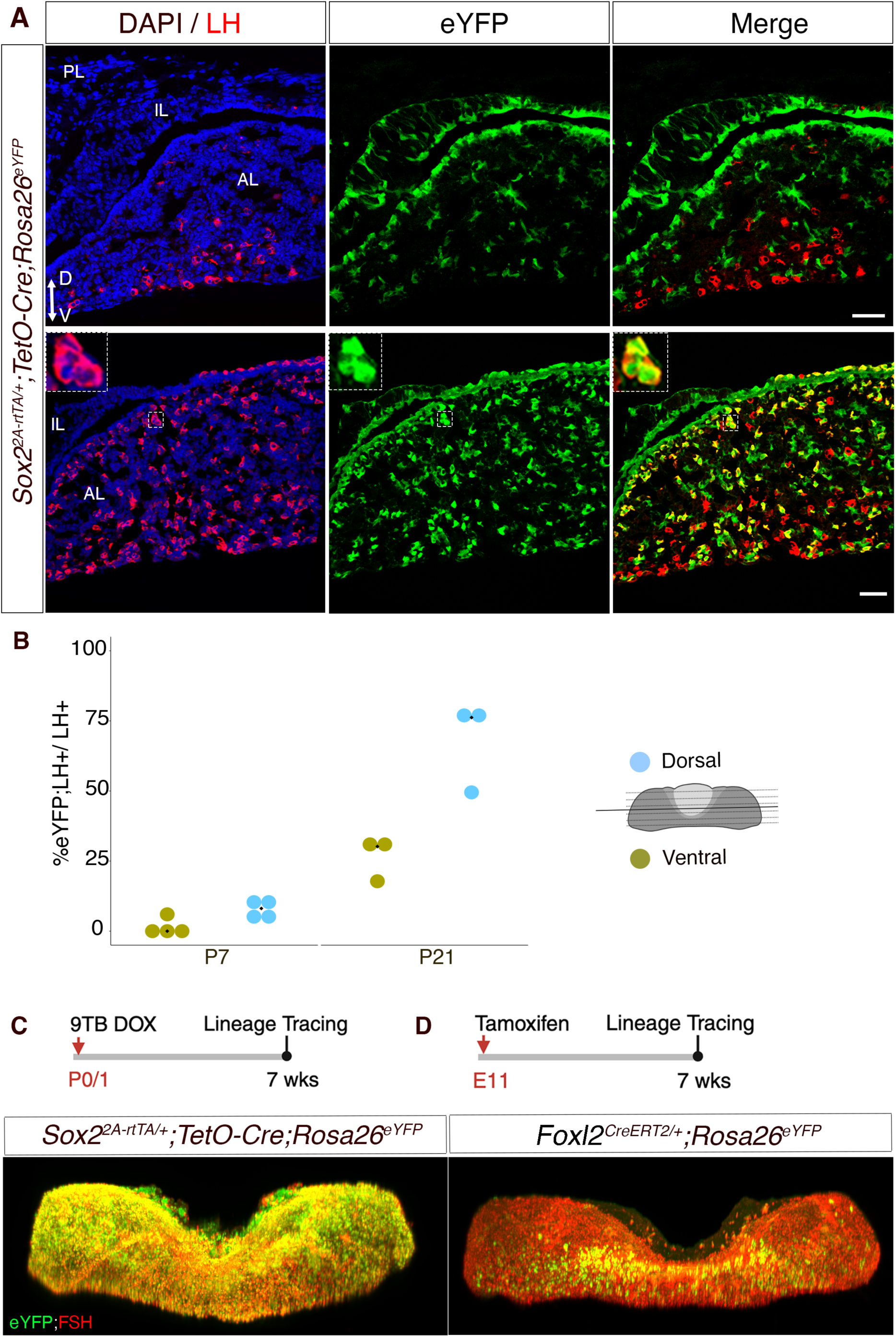
Postnatal and embryonic gonadotrophs occupy different domains in the pituitary. **A)** Immunofluorescent staining for LH and eYFP showing non-lineage traced ventral gonadotrophs at P7 (upper panel) in *Sox2rtTA;eYFP* pituitaries induced at birth. At P21 (lower panel), SC-derived lineage traced gonadotrophs are enriched dorsally. **B)** Quantification of differential localisation of SC-derived gonadotrophs. **C**, **D)** Whole-mount eYFP and FSH double immunofluorescence on a *Sox2rtTA;eYFP* pituitary induced at birth (C) and a *Foxl2CreERT2;eYFP* pituitary induced *in utero* (D), with the induction timing indicated above.

The presence of early eYFP^-ve^;LH^+ve^ cells in induced *Sox2rtTA;eYFP* pituitaries suggests that these cells have differentiated in the embryo (*4*). To assess this, we performed lineage tracing at 11.5dpc in *Foxl2^CreERT2/+^;Rosa26^ReYFP^*^/+^ embryos and performed whole mount immunofluorescence in adults, in parallel with *Sox2rtTa;eYFP* pituitaries induced at birth (Fig.4C). Despite a relatively poor induction efficiency with *Foxl2^CreERT2^*, we observe that eYFP;FSH double^+ve^ cells remain exclusively located near the medial-ventral surface of the anterior pituitary in contrast with the post-natal *Sox2rtTa;eYFP* gonadotrophs present throughout the gland.

In conclusion, neonate gonadotrophs emerge at least in part from cleft SCs and populate the entire gland with the exception of a small ventromedial domain exclusively populated by embryonic-born gonadotrophs, which persist into adulthood in both sexes.

### Differentiation of postnatal gonadotrophs is dictated by the physiological context, but is independent of GnRH and gonadal signals

To characterise mechanisms underlying predominant SC differentiation into gonadotrophs, and its temporal restriction to the early postnatal period, we first performed pituisphere assays to assess whether neonate SCs were primed toward gonadotroph fate acquisition. We differentiated in parallel neonate and adult pituispheres, as we previously described (*34*). We did not observe any preferential differentiation into gonadotrophs for postnatal spheres Instead, cells of both stages gave rise to LH and POU1F1 lineage cells in similar proportions (Fig.5A). This suggests that neonatal gonadotroph differentiation is dictated by context.

**Figure 5.**
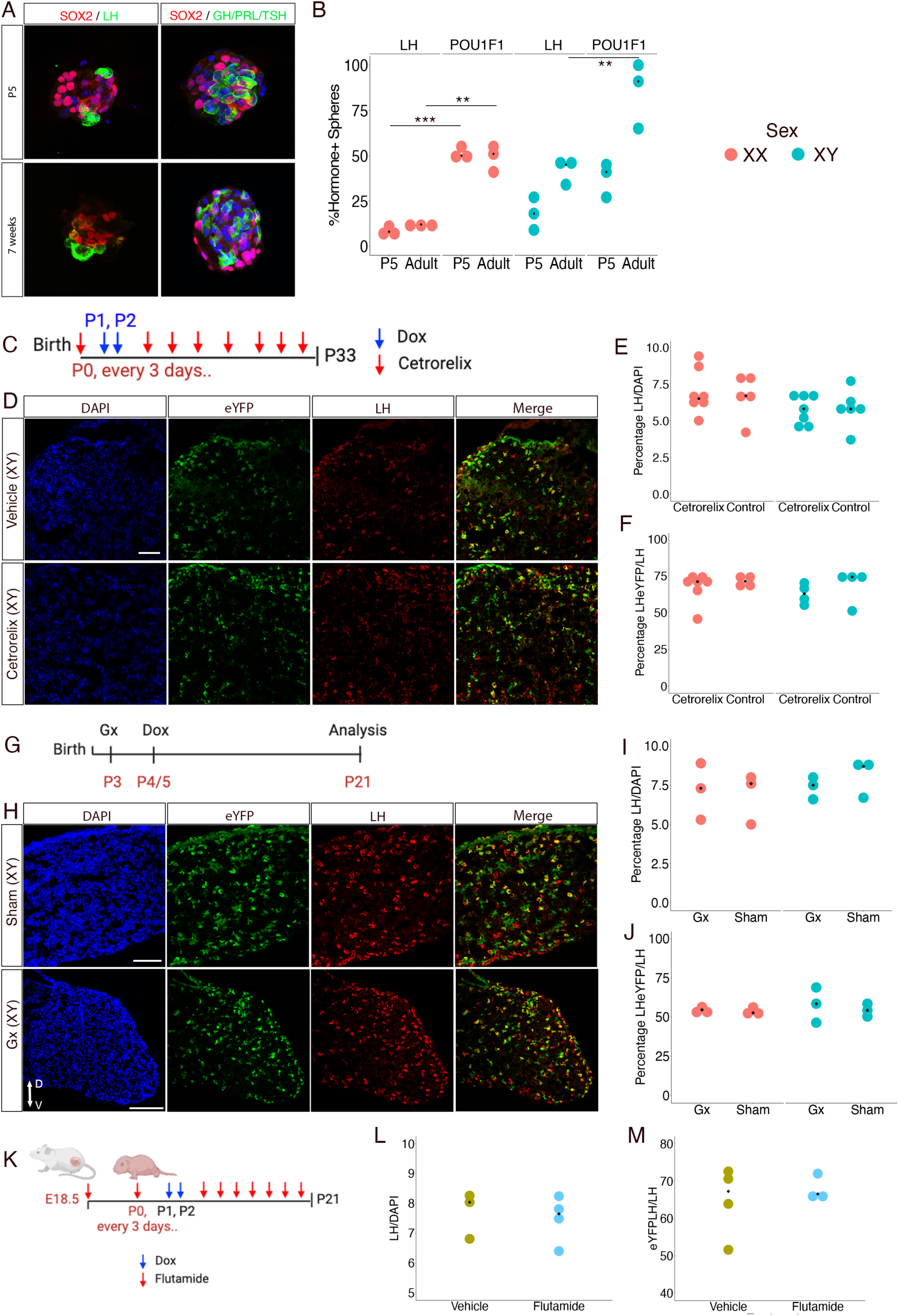
Regulation of neonatal SCs differentiation. **A**) Differentiation was induced in pituispheres from P5 (top) or 7 week (bottom) old male pituitaries. The presence of differentiated cells within was evaluated by immunofluorescence using LH or collectively GH, PRL and TSH antibodies for POU1F1 lineage cells detection. SOX2 staining confirms the presence of pituisphere forming-cells. **B**) The proportion of spheres containing LH or POU1F1 lineages cells were compared between ages in both sexes. Multiple unpaired-tests with Holm-Šídák post-hoc test. *P<0.05, **P<0.01, ***P<0.001, cultures from N=3 animals per sex and per age. **C**) Timeline of GnRH antagonist (cetrorelix) and lineage tracing induction (Dox) in *Sox2rtTa;eYFP* pups. **D**) Immunofluorescence for eYFP and LH in pituitaries of P33 cetrorelix treated male and control mice showing no apparent differences in SC-derived gonadotroph emergence. **E**-**F**) Quantification in both sexes at P33 shows no effect of cetrorelix on either the proportion of LH^+ve^/DAPI^+ve^ cells (E) or the proportion of eYFP; LH double^+ve^/LH^+ve^ cells (F). **G**) Timeline of gonadectomy and lineage tracing induction (Dox) in *Sox2rtTa;eYFP* pups. **H**) Immunofluorescence for eYFP and LH in pituitaries of P21 gonadectomised male and sham-operated control showing no apparent differences in SC-derived gonadotroph emergence. **I**-**J**) Quantification in both sexes at P21 shows no effect of gonadectomies on either the proportion of LH^+ve^/DAPI^+ve^ cells (I) or eYFP; LH double^+ve^/LH^+ve^ cells (J). **K**) Timeline of androgen antagonist (flutamide) treatment and lineage tracing induction (Dox) in *Sox2rtTa;eYFP* pups. **L**-**M**) Quantification in males at P21 shows no effect of flutamide on either the proportion of LH^+ve^/DAPI^+ve^ cells (L) or the proportion of eYFP; LH double^+ve^/LH^+ve^ cells (M). Scale bar = 80 μm for all images.

The abrupt cessation of maternal estrogen negative feedback at birth triggers pulsatile GnRH release, inducing minipubertal activation of the HPG axis (*1*). We thus examined a role for GnRH in inducing gonadotroph differentiation. SCs do not express its receptor, but we reasoned that embryonic gonadotrophs may relay a signal. We inhibited GnRH action by injecting the antagonist Cetrorelix (*35*) in *Sox2rtTa;eYFP* pups. Blockade of the HPG axis was monitored by examining gonads when pituitaries were harvested. We observed a clear gonadal size reduction in both sexes and a reduction in CYP17A1 staining in agreement with a downregulation of steroidogenesis in testes, consistent with reduced gonadotrophin levels following GnRH signalling blockade (Fig.S9). However, differentiation of SCs into gonadotrophs was not prevented and we observed a similar proportion of SC-derived gonadotrophs in treated and control animals (Figure 5C-F). We next investigated a role for gonadal steroids, because suppression of steroid feedback following pituitary target organ ablation affects pituitary adult SC activity (*9*). We performed gonadectomies in P3 *Sox2rtTa;eYFP* pups shortly before inducing lineage tracing at P4 and harvested pituitaries in 21-day-old animals. The proportion of eYFP;LH double^+ve^ gonadotrophs was not affected by gonad ablation (Figure 5H-J). In order to block effects of the neonatal male testosterone surge that occurs hours after birth, we used flutamide, an androgen-receptor antagonist (*36*). 18.5dpc pregnant dams and new-borns were dosed with flutamide, and treatment pursued every 3 days. Gonadotroph emergence at P21 was not affected (Figure 5K-M). Altogether, these results show that neonatal SC differentiation into gonadotrophs is influenced neither by GnRH, nor by gonadal signals. Despite impairment of the HPG axis in both experimental settings, SCs give rise to gonadotrophs in normal numbers and timing.

## Discussion

We demonstrate here that gonadotrophs have a dual origin, with the predominant proportion generated from resident SCs during minipuberty, and a smaller population of cells born in the embryo persisting in the adult gland. This heterogeneity is furthermore apparent in the gland with both populations occupying different domains: postnatal gonadotrophs emerge dorsally, in agreement with previous observations (*5-7*) and eventually invade the whole gland except for a small ventral region where cells specified in the embryo remain confined. This is the first evidence of the existence of two distinct subpopulations of gonadotrophs in the mouse. This phenomenon may extend to primates, because a sharp postnatal increase in gonadotroph numbers is observed in rhesus monkeys (*37*), suggesting its relevance to human physiology. While single cell transcriptomic analyses have so far failed to reveal subpopulations of gonadotrophs (*38*), heterogeneity in response to GnRH has been reported, although with an unknown basis (*39*). Furthermore, the intricacies of LH and FSH differential regulation in response to the same ligand, GnRH, are not yet completely understood, and manipulation of oestrogen hypothalamic feedback on the pulse generator hints at a further level of gonadotrophin regulation at the pituitary level (*40*). Thus, a dual origin and/or distinct localisation for gonadotrophs could explain aspects of their function and regulation. The existence of this dual population may also have implications for our understanding of a range of diseases affecting puberty, either delaying or preventing it, such as in patients affected by congenital hypogonadotropic hypogonadism (CHH), where GnRH signalling is impaired, or when it is induced precociously (*1*). For CHH patients, minipuberty has been suggested to represent a window of opportunity for early diagnoses and preventive treatment of pubertal disorders. Our findings give further support to the possibility of early postnatal intervention since this is when most gonadotrophs emerge.

Pituitary SCs can generate all pituitary endocrine cell types (*9, 10*). However, we show here that they predominantly generate gonadotrophs during the pituitary growth phase, encompassing minipuberty. Since neonate cells do not show this bias *in vitro*, this strongly suggests that the physiological context induces this event. We tested whether GnRH is involved since it initiates minipuberty, reasoning that embryonic gonadotrophs, the only pituitary cells expressing its receptor, may relay a signal to SCs. Antagonising GnRH signalling in neonates did not block gonadotroph emergence. This expands previous observations implying presence of gonadotrophs in absence of GnRH signalling. In CHH patients, fertility can indeed be restored by GnRH administration (*1*) and some *Gnrhr* mouse mutants can be rescued by pharmacological chaperones (*41*) demonstrating the presence of gonadotrophs. While GnRH signalling is involved during embryonic gonadotroph emergence (*4*), our results suggest that the gonadotrophs observed in animal models that lack GnRH signalling, contain this postnatal SC-derived population. We then tested whether gonadal feedback played a role by performing neonatal gonadectomies. In adults, removal of a pituitary target organ induces increased secretion, and sometimes numbers and differentiation from SCs of the cell type normally regulating the ablated organ (*9, 42*). However early postnatal gonadal ablation did not alter differentiation of neonate SCs into gonadotrophs in either sex, nor did perinatal onset of testosterone antagonism in males.

Gonadotrophs are the only pituitary endocrine population displaying such a clear dual origin. Since we did not observe a significant contribution of postnatal SCs to any other population, this implies that their expansion is explained by proliferation and/or differentiation of committed progenitors. It is worth highlighting that in addition to originating from embryonic versus postnatal SCs, which both come from Rathke’s pouch (RP) (*9*), other characteristics distinguish these SCs. RP cells express SOX2, but not SOX9, while postnatal SCs express both markers. It remains to be determined whether the properties of their gonadotroph progeny are influenced by the activity of these factors. While gonadotrophs may be unique in the pituitary with regard to their dual origin, other endocrine cells can show this distinction. Testicular Leydig cells first differentiate in the embryo, however this pool regresses postnatally to be mostly, but not completely, replaced by postnatal cells (*43*). Of interest, postnatal Leydig cells arise at the same time as the postnatal gonadotrophs. Indeed, postnatal gonadotrophs may induce this Leydig cell expansion because, while foetal Leydig cells develop independently of LH, postnatal cells require it.

Altogether these results show that GnRH and gonadal steroids, signals extrinsic to the pituitary that are well known for their action on gonadotrophs, are unlikely to play a role in differentiation of SCs into gonadotrophs. Perhaps it is the abrupt cessation of exposure to the maternal environment that may hold clues to the initiation of this postnatal wave of gonadotroph differentiation and thus mechanisms of minipuberty initiation. Finally, in addition to helping our understanding of congenital diseases affecting puberty, our findings may explain why the neonatal period is particularly sensitive to exposure to endocrine disruptors (*44*).

## Acknowledgments

We are grateful for the support from present and past members of the Lovell-Badge laboratory. We are indebted to the Francis Crick Institute platforms for their excellent support and expertise. We especially thank J. Mor from the Biological Research facility for taking excellent care of the animals, D. Bell from the Advanced Light microscopy, and experts from the Advanced Sequencing, Genetic modification service and Histology platforms.

## Author contributions

Conceptualization: K.R. and R.L-B. Methodology: D.S and K.R. Bioinformatic analyses: P.C and G.G. Investigation: D.S, K.R, Y.S, J.O, E.B and C.G. Visualization: D.S and K.R. Supervision: K.R and R.L-B. Writing—original draft: K.R and D.S. Writing—review and editing: R.L-B, Ph.M and P.M.

## Data and materials availability

All data needed to evaluate the conclusions in the paper are present in the paper and/or the Supplementary Materials. Datasets have been submitted in Gene Expression Omnibus (GSE275746), https://ncbi.nlm.nih.gov/geo/query/acc.cgi?acc=GSE275746.

## Material and Methods

### Mice

All animal experiments carried out were approved under the UK Animals (Scientific Procedures) Act 1986 and under the project licenses n. 80/2405 and PP8826065. *Sox2_tm1(RFP/rtTA)>rlb_* (this study), *Ednrb^em1(Ednrb-T2A-rtTA)Crick^* (this study), *GT(ROSA)26Sor^tm1(EYFP)Cos^ (45), (no gene)^Tg(tetO-Cre)LC1Bjd^(46), Lef1^tm1(EGFP/Cre/ERT2)Wtsi^* (*EMMA/* Infrafrontier, stock ID EM 10707) and *Foxl2^tm1(GFP/cre/ERT2)Pzg^* (JAX stock #015854) were maintained on C56BL/6Jax background.

### Generation of Sox2-T2A-TgRFP-T2A-rtTA mice

A Sox2-T2A-TgRFP-T2A-rtTA cassette was inserted in the *Sox2* locus by homologous recombination (GenOway) to generate *Sox2^tm1(RFP/rtTA)>rlb^* (Sox2^rtTA^). The neomycin resistance selection cassette was removed. While both copies of *Sox2* are retained, a proportion of animal homozygotes for the mutation appeared smaller, as observed in Sox2^+/-^ animals (*29*), suggesting that this mutation represents an hypomorphic allele for *Sox2*. Heterozygotes were normal and fertile and thus exclusively used in this study. *Sox2^rtTA/+^* were bred with *(no gene)^Tg(tetO-Cre)LC1Bjd^* and *GT(ROSA)26Sor^tm1(EYFP)Cos^*to perform lineage tracing experiments.

### Generation of Ednrb-T2A-rtTA mice

The *Ednrb*-T2A-rTA strain was generated by the Genetic Modification Service at the Francis Crick Institute using CRISPR-Cas9 assisted targeting to insert a dox-inducible reverse transactivator into exon 8 of the *Ednrb* gene via a GSG and T2A peptide linker and followed by a stop codon. The donor template contained a 774bp insert of GSG, T2A and rtTA3, with 692bp of homology at the 5’ end and 700bp homology at the 3’ end. The AAV donor vector was synthesised and packaged into AAV serotype 1 by VectorBuilder. The guide sequence used was 5’-ATAAATACAGCTCGTCTTGA-3’, that was synthesised as a synthetic guide RNA by IDT^TM^. AAV transduction and delivery of CRISPR Cas9 reagents were performed into 1-cell C57BL/6J zygotes as previously described (*47*).

### Cre Induction

For *Sox2^2A-rtTA/+^;TetO-Cre;Rosa26eYFP* and *Ednrb^2A-rtTA/+^;TetO-Cre;Rosa26eYFP* induction, two consecutive doses of 9-tert-Butyl Doxycycline HCl (9TBDox) (*30*) were administered by subcutaneous injection on two consecutive days between P0 and P4 (125µg/g body weight). For *Lef1^CreERT2/+^*;Rosa26^ReYFP/+^ pup induction, a single subcutaneous injection of tamoxifen (0.25 mg/g body weight) was administered. For *Foxl2^CreERT2/+^;Rosa26^ReYFP/+^* induction, tamoxifen (0.2 mg/g body weight) was administered to pregnant dam at 11.5 days post coitum and pups fostered at 19.5dpc.

### Cetrorelix treatment

Pups were administered with cetrorelix acetate (Sigma, C5249) diluted in water every 3 days, or water control, from P0 to P33 (3μg/g body weight) as described (*48*).

### Neonatal gonadectomies

Gonadectomies were performed at P3 in *Sox2^2A-rtTA/+^;TetO-Cre;Rosa26^eYFP^* neonates. In sham-operated animals, gonads were exposed.

### Flutamide treatment

Flutamide (Sigma-Aldrich, F9397) was dissolved in a solution of 10% ethanol in peanut oil to a dosage of 10 mg/ml/kg body weight. To block the effect of the neonatal surge of testosterone, pregnant dams were treated at 19.5 dpc. Treatment was initiated in pups at P0 and continued every three days until P21. Lineage tracing was induced with Dox at P1 and P2. The dosage was chosen based on reports of effective androgen receptor antagonism without significant secondary effects (such as changes in sex organ weight, gonadal development, anogenital distance, early-life body weight) or notable toxicity (*36*).

### Single-cell sequencing and immunofluorescence

Three *Sox9^iresGFP/+^*male and three *Sox9^iresGFP/+^* female P3 pituitaries were dissociated and cells sorted as described (*20*). Quality checks for single cell suspension and single cell sequencing were performed as described (*20*).Chromium Single Cell 30 Reagent Kits User Guide (v2 Chemistry) was used.

For immunofluorescence on plated cells, freshly dissociated cells were plated on Superfrost plus slides for 90 min in a cell incubator, fixed for 20 minutes on ice in 4% paraformaldehyde (PFA), and immediately processed for immunofluorescence.

### Bioinformatic analyses

Analyses were performed as previously described (*49*).

### Pituisphere culture

Dissociated anterior pituitary lobes (see above) were seeded at 50 × 10^3^ cells/ml in pituisphere medium containing EGF, FGF and 10% serum (*34*). After 7 days in culture, pituispheres were transferred into differentiating conditions (on Matrigel coated coverslips, without growth factor or serum) (*34*). Differentiated pituispheres were fixed for 30 minutes in 4% PFA at 4°C, and immunostaining was performed on the coverslips. Three independent repeats were performed.

### Immunofluorescence, analyses of cell proliferation and RIA

Pituitaries were fixed overnight in 4% PFA at 4°C. Immunofluorescent stainings were performed on cryosections as previously described (*49*). The following primary antibodies were used: Goat anti-SOX2 (Biotechne, AF2018), rabbit anti-SOX9 (a gift from F. Poulat, IGH Montpellier, France), rat anti-GFP (Nacalai Tesque, 04404-84), rabbit anti-Lef1(Abcam, ab137872), and hormone antibodies anti-LH, GH, POMC, and PRL from the National Hormone Peptide Program (A.F. Parlow, Torrance, USA). Slides were then incubated with Alexa Fluor secondary antibodies. Imaging of stained tissue sections was performed on a Leica SPE microscope and imaging of plated cells was done on an Olympus spinning disk microscope.

For proliferation analyses, pituitaries were harvested following a one-hour EdU pulse (30μg/g body weight). EdU incorporation was detected using the Click-iT EdU imaging kit following the manufacturer instructions (Thermo Fisher Scientific). Sections were washed with PBST and mounted using Aqua-Poly/Mount (Polysciences, Inc., Warrington, PA, USA).

Radioimmunoassays (RIA) were performed as described (*49*).

### Whole mount immunofluorescence

The iDISCO+ protocol was followed, as described (https://idisco.info/wp-content/uploads/2015/04/whole-mount-staining-bench-protocol-methanol-dec-2016.pdf) using Chick anti-GFP (Aves, GFP-1020) and Rabbit anti-FSH from the National Hormone Peptide Program (A.F. Parlow, Torrance, USA). Staining penetration was improved by extending primary antibody incubation time up to 2 weeks. Clearing was performed in a benzyl alcohol/benzyl benzoate 1:2 solution. Samples were then transferred and kept in ethyl cinnamate. Imaging was performed on a Light Sheet Lavision UM II microscope and images reconstructed using the Imaris software.

### Cell Counts

For dissociated cell counts, slides stained by immunofluorescence were scanned. Automated counts were then performed using QuPath-0.3.2 (https://qupath.github.io/). We routinely counted between 100,000 cells at P5 to 300,000 cells in adults.

SOX2^+ve^ cells were counted manually on sections (Fig.1C) using Fiji because the staining on dissociated cells was not satisfactory. For *Sox2^2A-rtTA/+^;TetO-Cre;Rosa26eYFP;* or *Lef1^CreERT2/+^*;Rosa26^ReYFP/+^ eYFP;hormone double^+ve^ cell counting, pituitaries were serially sectioned on 5 slides. One slide was immunostained for eYFP and the chosen hormonal marker and all double^+ve^ cells manually identified. Manually identified double^+ve^ cells were individually imaged on a SPE Leica confocal microscope and images subsequently reviewed to assess co-localisation, and double^+ve^ cells scored.

All counting results and tests are presented in supplementary tables S3 and S4.

