## Supplementary figures for "Gonadotrophs have a dual origin, with most derived from pituitary stem cells during minipuberty"

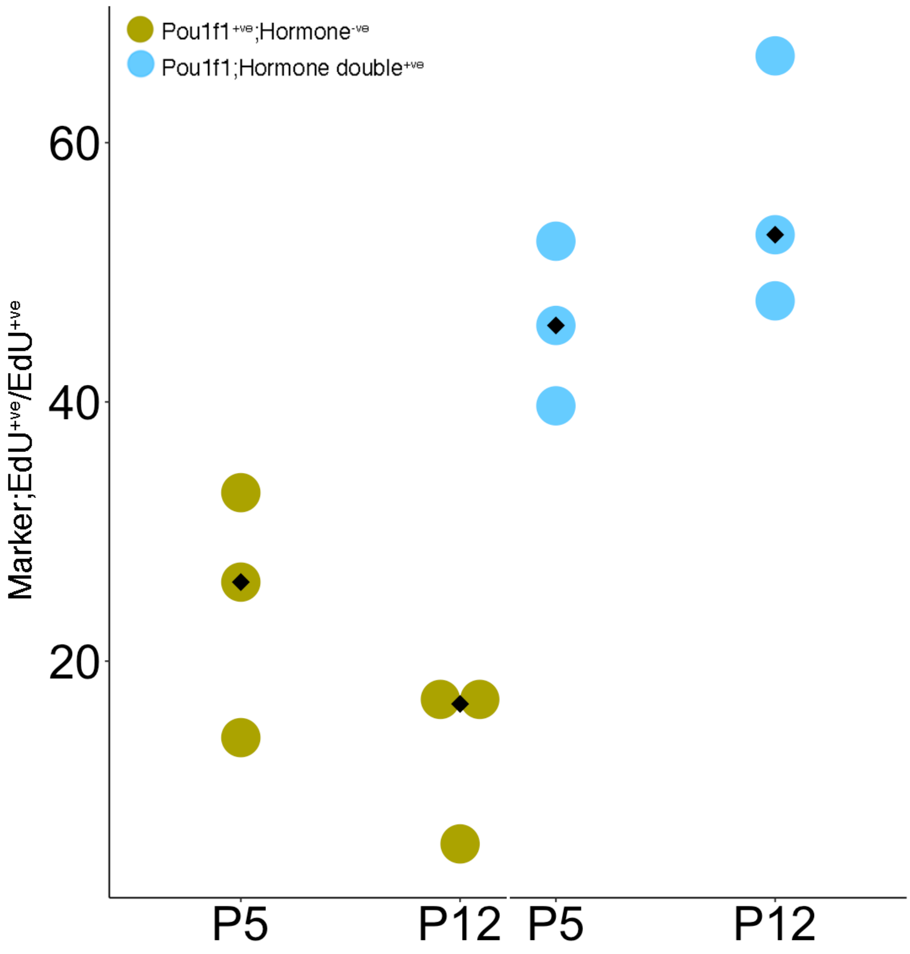


**Supplementary Fig.1: Quantification of POU1F1$^{+ve}$ cells proliferation.**

Proliferation was assessed by quantifying the number of EdU$^{+ve}$ cells in male mice pituitaries after a one-hour pulse at the ages indicated. Quantification was performed manually on sections stained by immunofluorescence for POU1F1 and mix of rabbit antibodies recognising all POU1F1 lineage endocrine cell types (GH, Prl, TSH). A comparable percentage of hormone$^{+ve}$ and hormone$^{-ve}$ POU1F1$^{+ve}$ cells proliferate at P5 and P12 with no significant difference between timepoints or cell type. Each dot represents one animal.

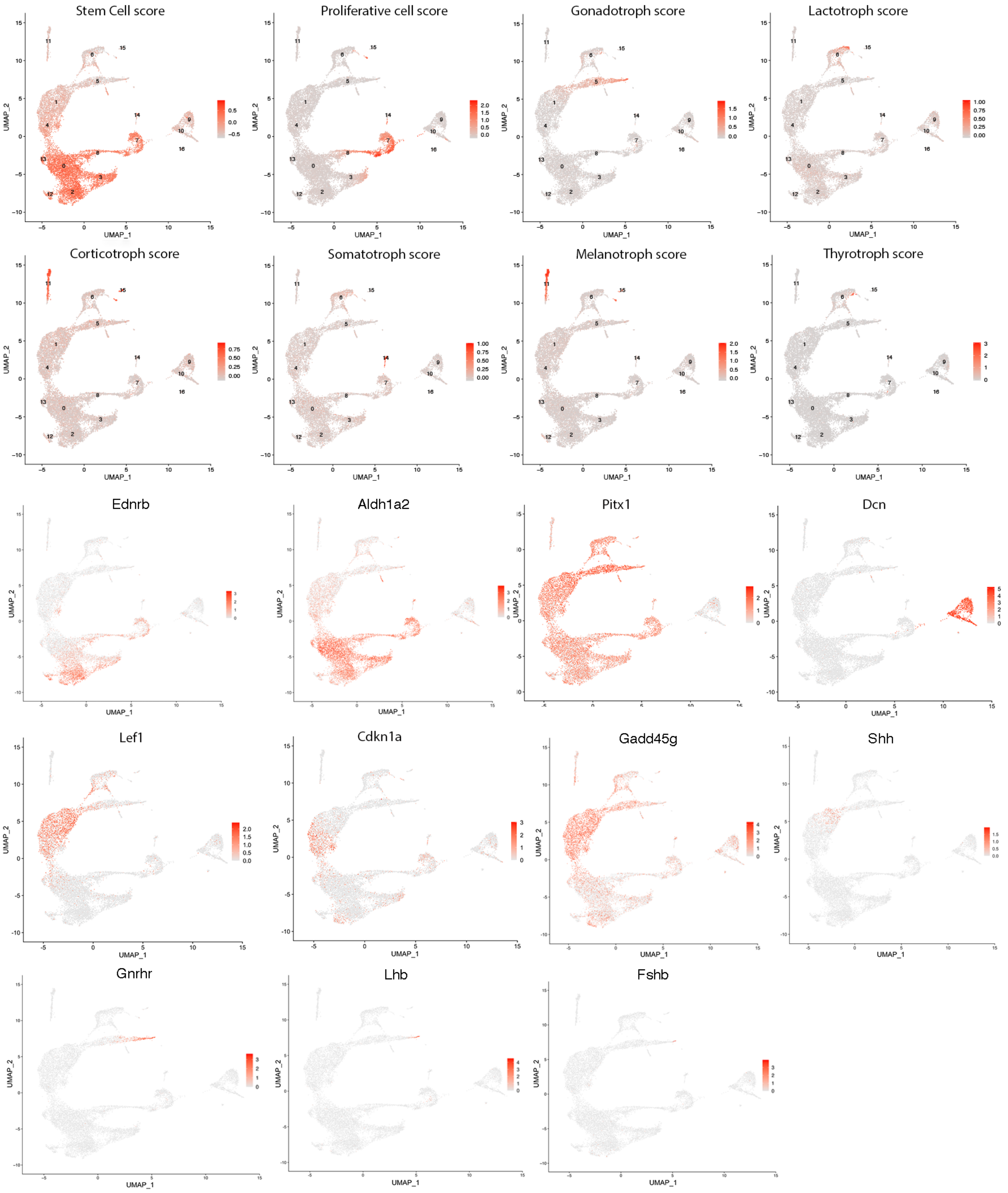


**<u>Supplementary Fig2</u>: UMAP representation of cell type scores and selected marker expression.** We used the same cell type signatures as previously (*20*).

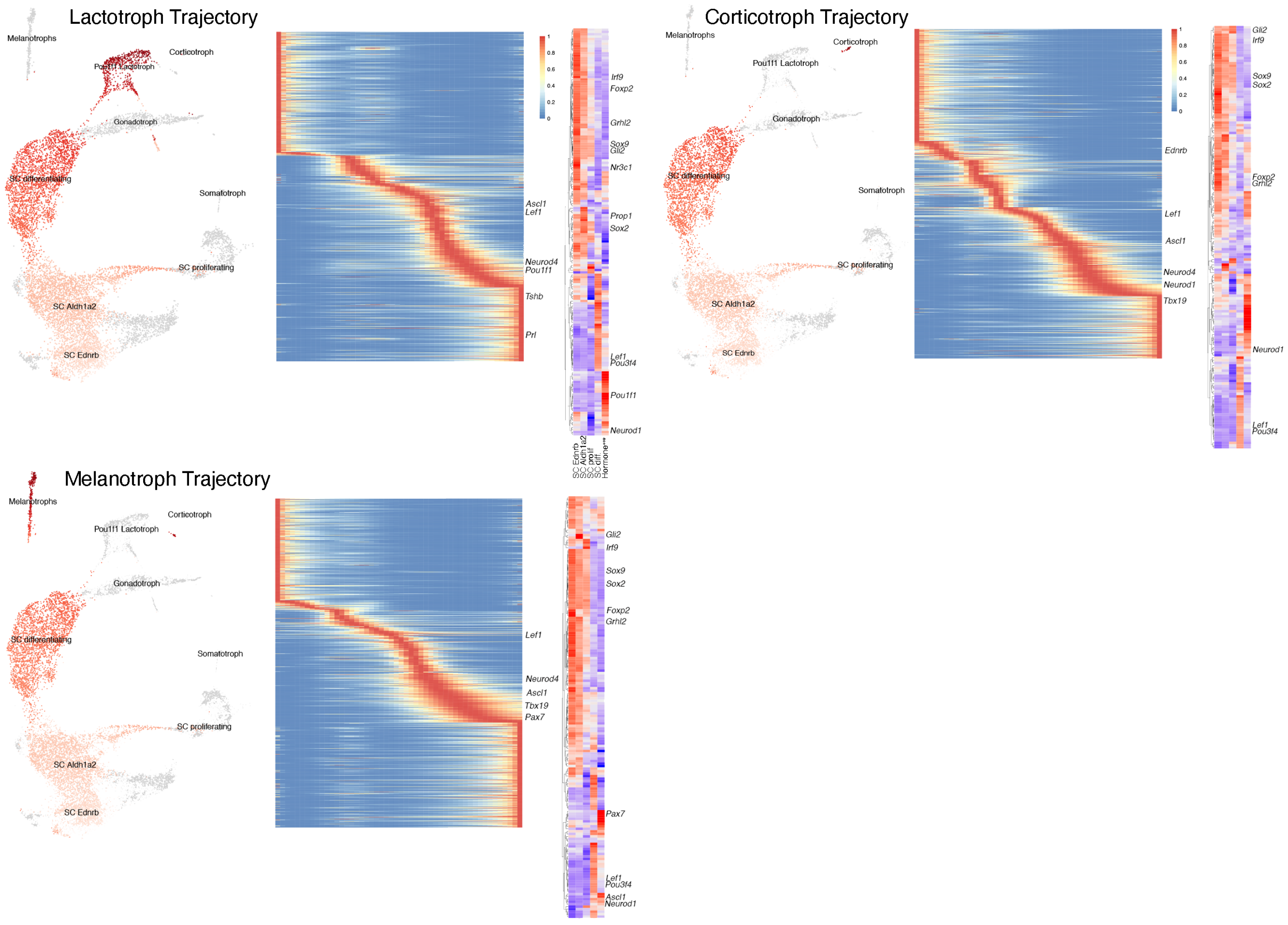


**<u>Supplementary Fig.3</u>: Slingshot trajectories and heatmaps for lactotroph, corticotroph and melanotroph trajectories.**

Genes common to all trajectories are highlighted. The trajectories presented include endocrine lineages to which SCs contribute, as demonstrated by lineage tracing analyses (Fig.3).

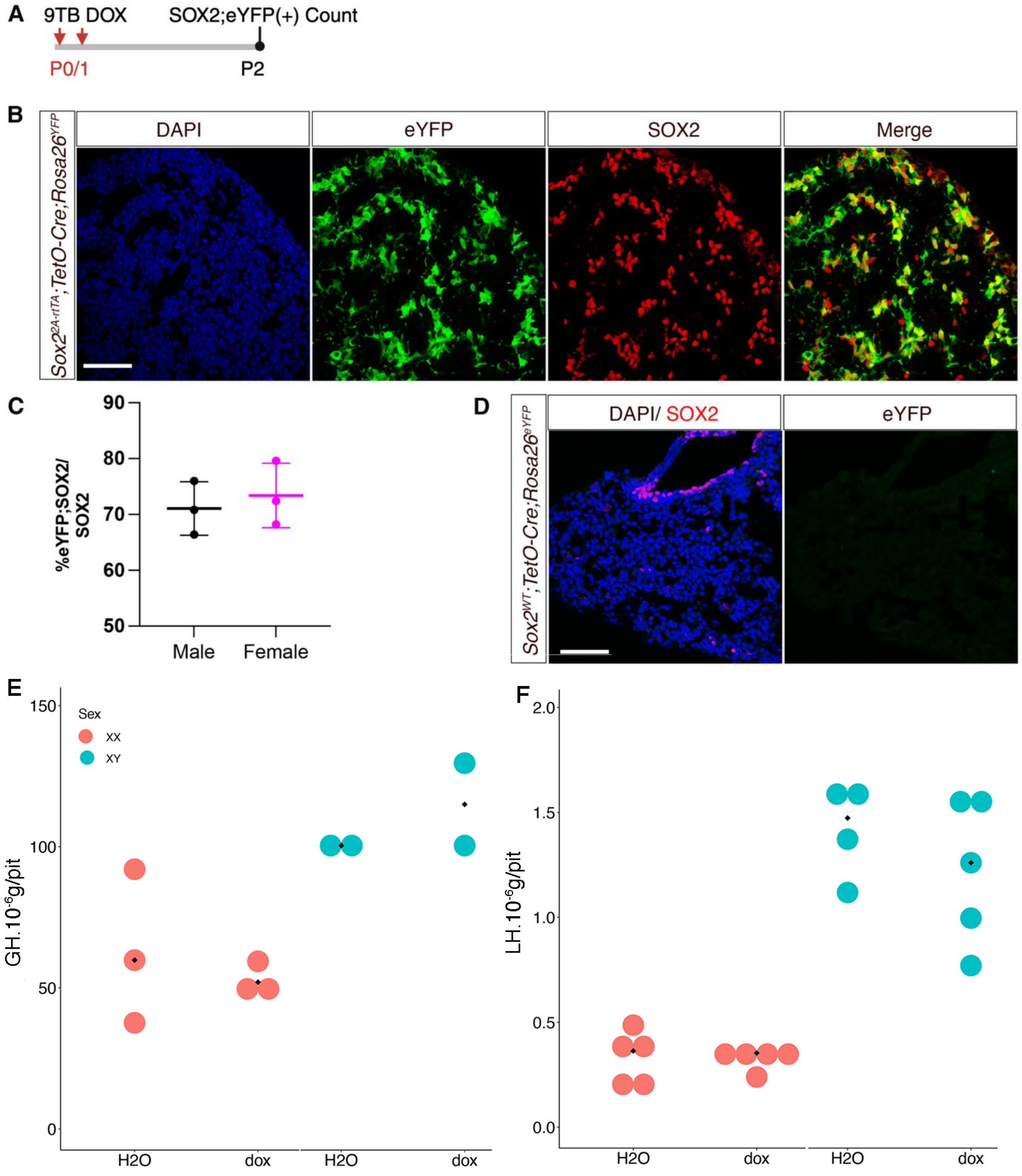


**<u>Supplementary Fig.4: Characterisation of the Sox2rtTA allele.</u>**

**A**) Timeline for *Sox2rtTA;eYFP* efficiency assessment. 9TB-Dox was administered at P0 and P1 and the percentage of SOX2;eYFP double$^{+ve}$ cells counted at P2. **B**) Immunostaining of eYFP and SOX2 shows most SOX2$^{+ve}$ cells express eYFP. **C**) Percentage of eYFP;SOX2 double$^{+ve}$ cells as a proportion of the total number of SOX2$^{+ve}$ cells; recombination efficiency is approximately 70% (N=3 per sex). **D**) No expression of eYFP is detected in the absence of the *Sox2rtTA* allele. **E**,**F**) 9TB-Dox or vehicle (water) was injected on two consecutive days between P0 and P4. Pituitaries were harvested at 7 weeks and GH (E) and LH (F) levels measured by RIA . There was no significant difference in LH and GH levels between treated and control animals (Mann-Whitney test, GH $p = 0.67$ for males and 0.70 for females LH $p = 0.41$ for males and 0.84 for females). Each dot represents one animal. Scale bars represent 50 μm.

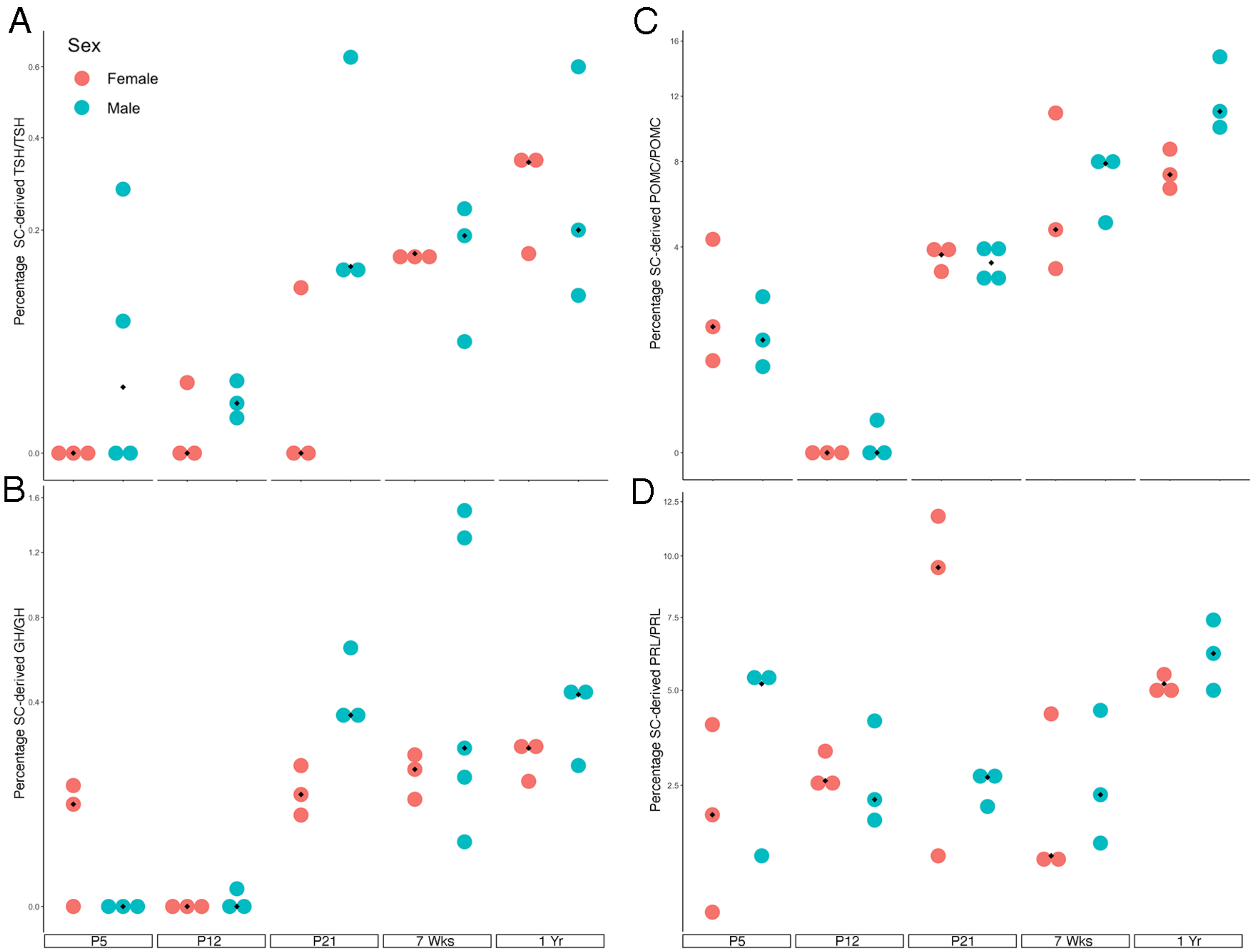


**<u>Supplementary Fig.5:</u> Timeline of SC-derived endocrine cell emergence.**
Beside gonadotrophs (Fig.3) SCs mostly contribute to POMC lineage (corticotrophs and melanotrophs, **C**), and lactotrophs (PRL, **D**). There is very little contribution to somatotrophs (GH, **A**) and thyrotrophs (TSH, **B**). Each dot represents one animal.

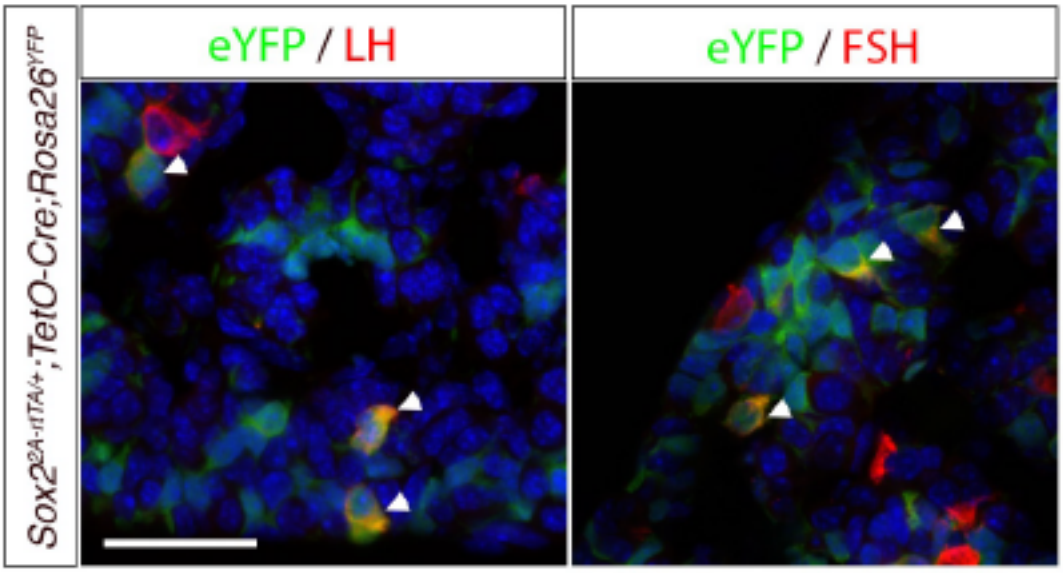


**<u>Supplementary Fig.6:</u> Newly differentiated gonadotrophs express both LH and FSH.** Double immunofluorescences for eYFP and LH and eYFP and FSH in a P4 female pituitary. 9TBD induction was performed at P0 and P1. Some cells in the progeny of $SOX2^{+ve}$ SC have already acquired a gonadotroph fate with both LH and FSH being upregulated. The scale bar represents 30µm.

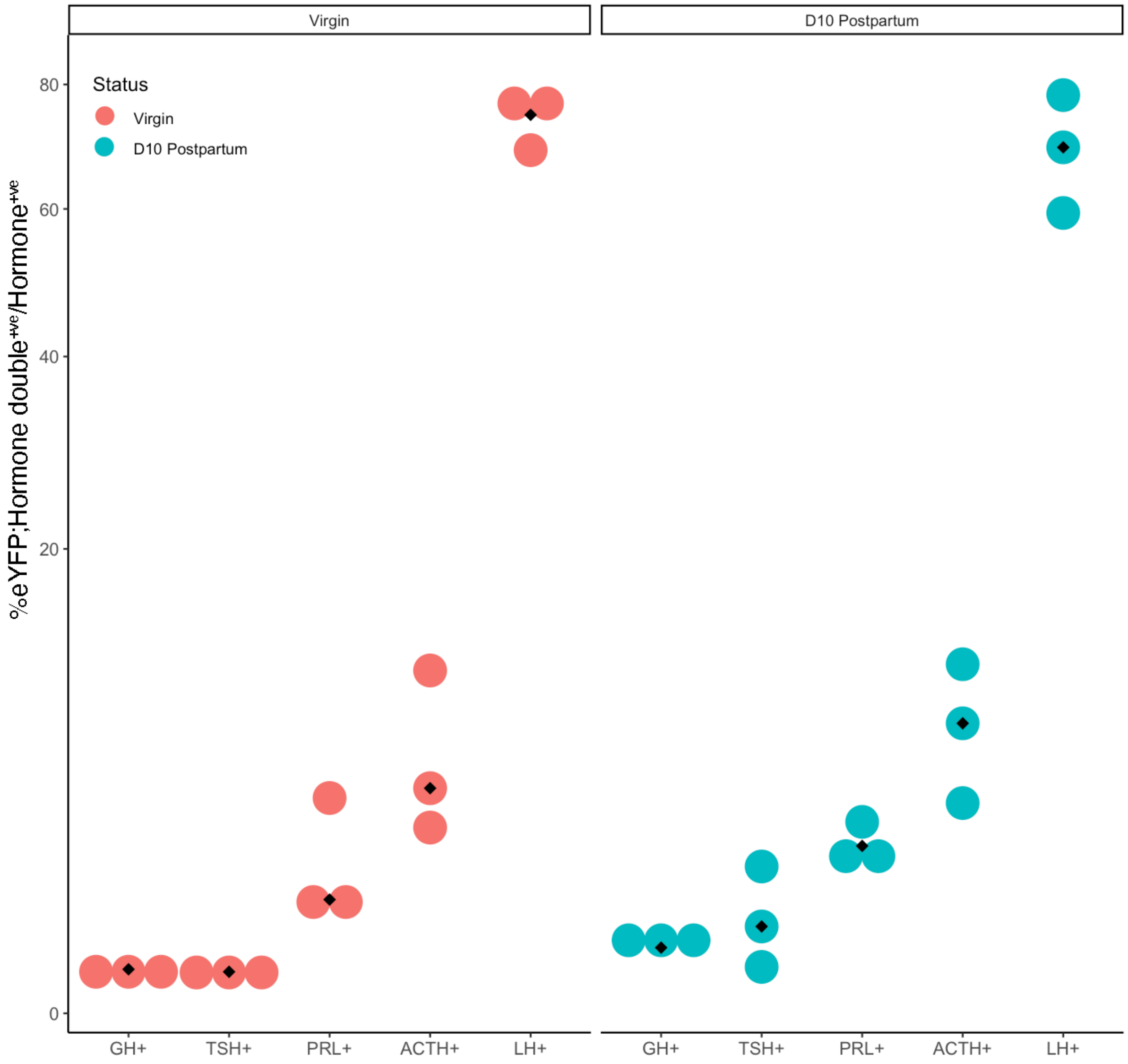


**Supplementary Fig.7: Effect of pregnancy and lactation on female pituitary SC contribution on day 10 postpartum.**

Lineage tracing of *Sox2rtTA;eYFP* female pups was initiated at P0/P1 and animals mated at 7 weeks. Following their first litter at approximately 10 weeks, pituitaries were harvested from lactating dams on day 10 postpartum and the percentage of eYFP;hormone double$^{+ve}$ cells counted for each cell type. Multiple unpaired t-tests were performed between virgin and lactating mice, and the Holm-Šídák method used to correct for multiple comparisons. No significant difference in SC contribution was observed between virgin and lactating females.

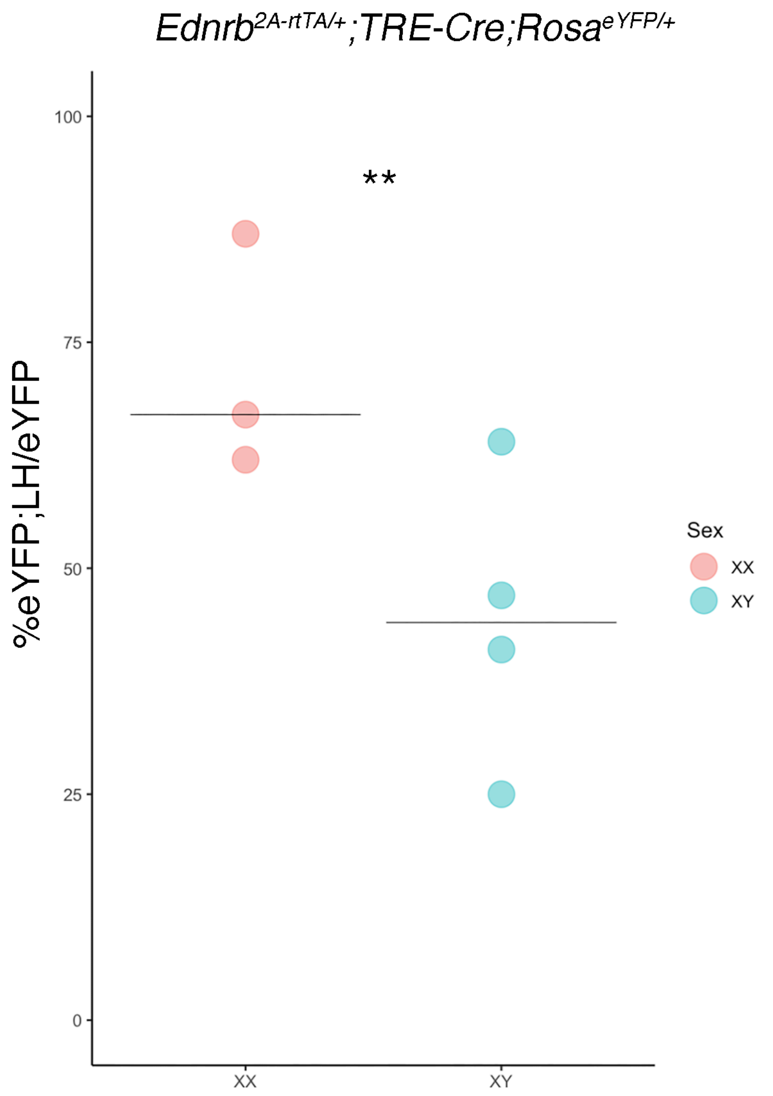


<u>**Supplementary Fig.8**</u>**: Cleft SC lineage tracing in *Ednrb$^{2A\text{-}rtTA/+}$;TetO-Cre;Rosa26$^{ReYFP/+}$* animals.**

Pups were treated at P0 and P1 using 9TB-Dox, in the same conditions as *Sox2rtA;eYFP* animals and pituitaries harvested at or after 7 weeks. Tracing was relatively inefficient, probably reflecting the low levels of EDNRB observed *in vivo* (*20*). Traced endothelial cells, morphologically recognisable, were not included in the analysis. In agreement with the data obtained in Sox2rtTA;eYFP analyses, gonadotrophs were mostly represented in EDNRB progeny. Beta regression was used to assess the effect of sex on proportion of LH positive cells in the endocrine progeny of EDNRB$^{+ve}$ cells ($p= 0.007629385$) and showed that there are more LH$^{+ve}$ cells in the progeny of female cleft SCs. Each dot represents one animal.

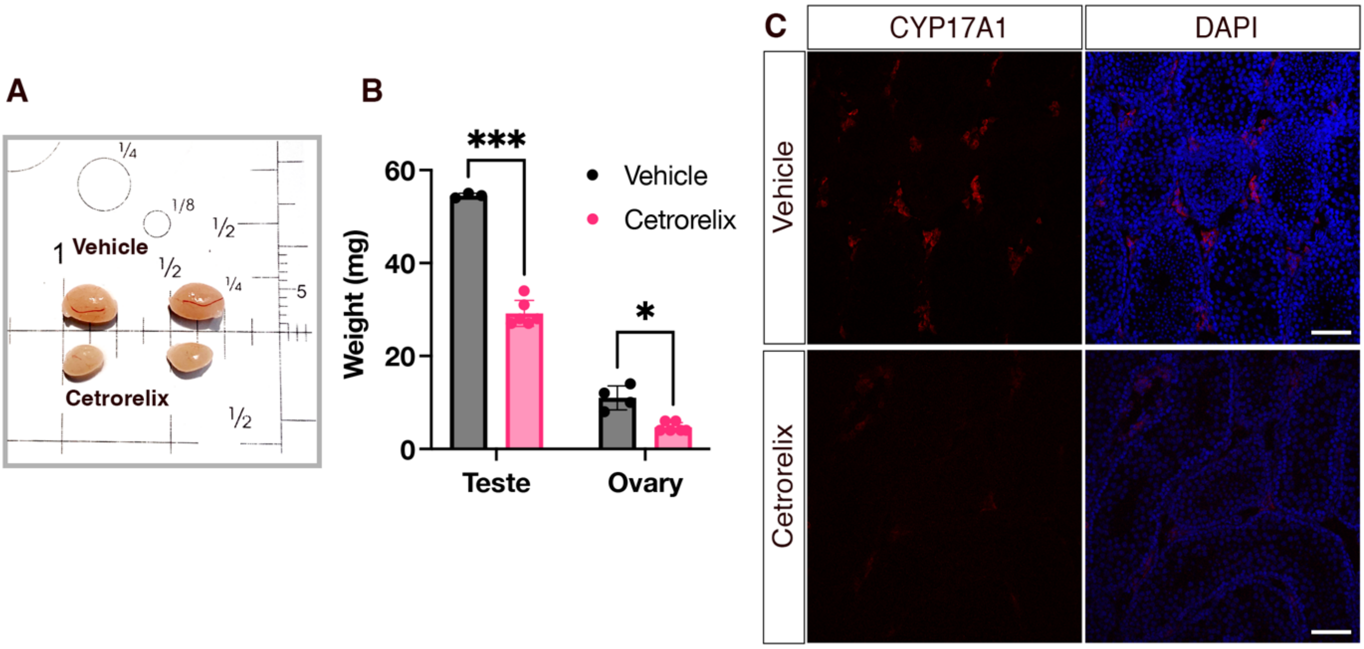


**<u>Supplementary Fig.9</u>: Cetrorelix-induced gonadal hypomorphism.**

**A**) Reduction of testes size in P33 cetrorelix treated animals comparable to vehicle controls. **B**) Both testes and ovaries of treated animals show a reduction in weight. **C**) CYP17A1 immunofluorescence on testis section shows reduction in cetrorelix treated animals demonstrating reduction in Leydig cells steroidogenesis, induced by lack of gonadotrophins.

**Sup. Table 1. Percentage of endocrine cells in the maturing postnatal pituitary.**

Percentage, standard deviation and mean for each time point, calculated as the percentage of Hormone;DAPI double $^{+ve}$/DAPI, following removal of the posterior and intermediate lobes (graph shown Fig1A-B). Multiple unpaired t-tests (Mann-Whitney Rank test) were performed between sexes for each cell type at each time point, and the two-stage step-up (Benjamini, Krieger, and Yekutieli) method used to correct for multiple comparisons.

| Age | Sex | Somatotroph | | | | | Lactotroph | | | | | Corticotroph | | | | |
|---|---|---|---|---|---|---|---|---|---|---|---|---|---|---|---|---|
| | | % | STD | q Val | FC P5 | STD | % | STD | q Val | FC | STD | % | STD | q Val | FC P5 | STD |
| P5 | XX | 13.34 | 2.77 | 0.272 | 1.00 | 0.21 | 0.91 | 0.62 | 0.424 | 1.00 | 0.68 | 8.01 | 2.92 | 0.03 | 1.00 | 0.36 |
| | XY | 10.70 | 1.06 | | 1.00 | 0.10 | 0.59 | 0.37 | | 1.00 | 0.63 | 11.65 | 1.54 | | 1.00 | 0.13 |
| P12 | XX | 8.47 | 1.41 | 0.398 | 0.63 | 0.11 | 11.70 | 2.26 | 0.639 | 12.86 | 2.48 | 9.92 | 2.05 | 0.93 | 1.24 | 0.26 |
| | XY | 13.18 | 6.18 | | 1.23 | 0.58 | 10.77 | 2.14 | | 18.25 | 3.62 | 9.80 | 2.17 | | 0.84 | 0.19 |
| P21 | XX | 37.16 | 4.28 | 0.425 | 2.79 | 0.32 | 13.83 | 2.73 | 0.451 | 15.20 | 3.00 | 11.94 | 1.69 | **0.02** | 1.49 | 0.21 |
| | XY | 35.36 | 4.00 | | 3.30 | 0.37 | 12.58 | 3.24 | | 21.32 | 5.49 | 8.40 | 2.44 | | 0.72 | 0.21 |
| 7 wks | XX | 23.75 | 3.03 | **0.031** | 1.78 | 0.23 | 34.80 | 5.35 | **0.025** | 38.24 | 5.88 | 7.15 | 0.92 | 0.25 | 0.89 | 0.11 |
| | XY | 37.40 | 6.84 | | 3.50 | 0.64 | 25.06 | 1.95 | | 42.47 | 3.30 | 8.39 | 1.43 | | 0.72 | 0.12 |
| 1 year | XX | 23.25 | 8.25 | 0.328 | 1.74 | 0.62 | 43.21 | 6.06 | **0.025** | 47.49 | 6.66 | 6.07 | 0.65 | 0.93 | 0.76 | 0.08 |
| | XY | 28.59 | 5.04 | | 2.67 | 0.47 | 25.93 | 9.45 | | 43.95 | 16.02 | 6.26 | 1.91 | | 0.54 | 0.16 |
| | | Gonadotroph | | | | | Thyrotroph | | | | | | | | | |
| P5 | XX | 2.00 | 0.26 | 0.543 | 1.00 | 0.13 | 4.91 | 0.50 | 0.224 | 1.00 | 0.10 | | | | | |
| | XY | 1.57 | 0.54 | | 1.00 | 0.34 | 5.42 | 0.66 | | 1.00 | 0.12 | | | | | |
| P12 | XX | 1.97 | 0.70 | >0.999 | 0.98 | 0.35 | 3.35 | 0.73 | 0.224 | 0.68 | 0.15 | | | | | |
| | XY | 1.79 | 0.21 | | 1.14 | 0.13 | 4.06 | 0.52 | | 0.75 | 0.10 | | | | | |
| P21 | XX | 5.90 | 0.61 | >0.999 | 2.95 | 0.31 | 1.50 | 0.43 | **0.041** | 0.31 | 0.09 | | | | | |
| | XY | 6.05 | 1.66 | | 3.86 | 1.06 | 2.44 | 0.55 | | 0.45 | 0.10 | | | | | |
| 7 wks | XX | 7.42 | 3.24 | >0.999 | 3.71 | 1.62 | 1.59 | 0.19 | 0.224 | 0.32 | 0.04 | | | | | |
| | XY | 7.36 | 3.28 | | 4.69 | 2.09 | 2.08 | 0.63 | | 0.38 | 0.12 | | | | | |
| 1 year | XX | 5.02 | 1.51 | 0.083 | 2.51 | 0.76 | 1.16 | 0.42 | 0.550 | 0.24 | 0.08 | | | | | |
| | XY | 2.92 | 0.91 | | 1.86 | 0.58 | 1.02 | 0.22 | | 0.19 | 0.04 | | | | | |

**Sup. Table 2. Percentage of proliferating endocrine cells in the postnatal pituitary.**

Mean percentage and standard deviation reported for each time point, calculated as the percentage of Hormone;EdUdouble$^{+ve}$/Hormone$^{+ve}$, following removal of the posterior and intermediate lobes (Graph shown Fig.1C). To establish sex differences in proliferation rates across time points, multiple unpaired t-tests were performed and the Holm-Šídák method was used to correct for multiple comparisons.

| Age | Sex | GH | | | PRL | | | POMC | | |
|---|---|---|---|---|---|---|---|---|---|---|
| | | % | STD | Adj P Val | % | STD | Adj P Val | % | STD | Adj P Val |
| P5 | XX | 5.87 | 1.76 | 0.792302 | 3.89 | 1.47 | 0.514020 | 4.50 | 1.40 | 0.758315 |

| | | | | | | | | | | |
|---|---|---|---|---|---|---|---|---|---|---|
| | XY | 4.77 | 2.57 | | 6.35 | 2.35 | | 3.83 | 0.21 | |
| **P12** | XX | 3.43 | 0.12 | 0.419445 | 5.77 | 1.07 | 0.576953 | 2.63 | 0.58 | 0.749082 |
| | XY | 3.07 | 0.31 | | 5.04 | 0.35 | | 3.05 | 0.36 | |
| **P21** | XX | 1.00 | 0.44 | 0.827545 | 3.00 | 0.89 | 0.576953 | 0.27 | 0.15 | 0.889507 |
| | XY | 0.88 | 0.16 | | 3.63 | 0.51 | | 0.29 | 0.06 | |

| **Age** | **Sex** | **LH** | | | **TSH** | | | **SOX2** | | |
|---|---|---|---|---|---|---|---|---|---|---|
| | | **%** | **STD** | **Adj P Val** | **%** | **STD** | **Adj P Val** | **%** | **STD** | **Adj P Val** |
| **P5** | XX | 1.03 | 0.21 | 0.11 | 3.07 | 0.21 | 0.10 | 8.20 | 0.53 | 0.04 |
| | XY | 0.53 | 0.25 | | 2.03 | 0.47 | | 17.23 | 2.80 | |
| **P12** | XX | 0.44 | 0.13 | 0.14 | 0.41 | 0.14 | 0.17 | 8.87 | 0.95 | 0.002 |
| | XY | 0.63 | 0.15 | | 0.70 | 0.20 | | 2.63 | 0.55 | |
| **P21** | XX | 0.00 | 0.00 | N/A | 0.90 | 0.87 | 0.17 | 0.80 | 0.15 | 0.51 |
| | XY | 0.00 | 0.00 | | 0.11 | 0.12 | | 0.73 | 1.27 | |

**Sup. Table 3 Percentage of endocrine cells at 7 weeks of age generated postnatally from SOX2$^{+ve}$ SCs.**

Percentage of eYFP;Hormone double$^{+ve}$/Hormone in *Sox2rtTA;eYFP*. No significant difference was observed between sexes in mean SC-contribution for any lineage. Multiple unpaired t-tests were performed and the Holm-Šídák method was used to correct for multiple comparisons.

| **Cell Type** | **XX** | | **XY** | | |
|---|---|---|---|---|---|
| | **Mean (%)** | **STD** | **Mean (%)** | **STD** | **Adj P val** |
| **GH** | 0.17 | 0.06 | 0.65 | 0.69 | 0.966 |
| **TSH** | 0.16 | 0.01 | 0.16 | 0.10 | 0.966 |
| **PRL** | 2.20 | 1.82 | 2.70 | 1.54 | 0.922 |
| **POMC** | 6.27 | 4.08 | 7.00 | 1.73 | 0.966 |
| **LH** | 74.93 | 5.03 | 71.61 | 10.54 | 0.966 |

**Sup. Table 4 Temporal lineage tracing of SC contribution to all lineages.**

Percentage of eYFP;Hormone double$^{+ve}$/Hormone in *Sox2rtTA;eYFP*. No significant difference was observed in mean %eYFP-positivity between sexes. Multiple unpaired t-tests were performed and the Holm-Šídák method was used to correct for multiple comparisons.

| | Age | XX | | XY | | |
|---|---|---|---|---|---|---|
| | | Mean (%) | STD | Mean (%) | STD | Adj. P Value |
| LH+ | P5 | 10.01 | 2.97 | 9.8 | 0.91 | 0.935 |
| | P12 | 43.51 | 9.12 | 45.88 | 1.29 | 0.935 |
| | P21 | 56.31 | 0.5 | 51.94 | 6.8 | 0.790 |
| | 7 weeks | 74.93 | 5.03 | 76.16 | 4.97 | 0.935 |
| | 1 year | 76.16 | 2.49 | 69.16 | 3.63 | 0.232 |
| PRL+ | P5 | 2.13 | 1.76 | 4.00 | 2.43 | 0.791 |
| | P12 | 2.80 | 0.44 | 2.70 | 1.23 | 0.896 |
| | P21 | 7.50 | 5.58 | 2.49 | 0.37 | 0.791 |
| | 7 weeks | 2.20 | 1.82 | 2.70 | 1.54 | 0.896 |
| | 1 year | 5.17 | 0.35 | 6.20 | 1.20 | 0.791 |
| GH+ | P5 | 0.08 | 0.07 | 0.00 | 0.00 | 0.562 |
| | P12 | 0.00 | 0.00 | 0.00 | 0.00 | 0.571 |
| | P21 | 0.13 | 0.06 | 0.45 | 0.17 | 0.130 |
| | 7 weeks | 0.17 | 0.06 | 0.65 | 0.69 | 0.571 |
| | 1 year | 0.21 | 0.06 | 1.08 | 0.57 | 0.571 |
| TSH+ | P5 | 0.00 | 0.00 | 0.09 | 0.13 | 0.621 |
| | P12 | 0.01 | 0.01 | 0.01 | 0.01 | 0.725 |
| | P21 | 0.04 | 0.06 | 0.10 | 0.07 | 0.394 |
| | 7 weeks | 0.16 | 0.01 | 0.31 | 0.30 | 0.988 |
| | 1 year | 0.28 | 0.11 | 0.30 | 0.27 | 0.988 |
| POMC+ | P5 | 2.20 | 1.85 | 1.40 | 0.82 | 0.925 |
| | P12 | 0.00 | 0.00 | 0.03 | 0.06 | 0.889 |
| | P21 | 3.63 | 0.50 | 3.41 | 0.65 | 0.925 |
| | 7 weeks | 6.27 | 4.08 | 7.00 | 1.73 | 0.925 |
| | 1 year | 7.53 | 1.07 | 11.93 | 2.53 | 0.263 |
